# Glycoconjugates tune the viscoelastic signature of mammary tumor progression

**DOI:** 10.64898/2026.10.20.760544

**Authors:** Abhishek Sharma, Jane A. Baude, Maya Papez, Marissa A. Gionet-Gonzales, Ryan S. Stowers

## Abstract

Breast cancer progression involves extensive remodeling of the tumor extracellular matrix (ECM), resulting in changes in mechanical properties. Tumors are viscoelastic, exhibiting both an initial elastic response to loading and a time-dependent viscous response. While it is well established that tumor elasticity or stiffness is elevated, changes in tissue viscoelasticity during breast cancer progression remain poorly characterized. Here, using a mouse model that recapitulates stages of human breast cancer progression (mouse mammary tumor virus-polyoma middle T antigen, MMTV-PyMT), we show that tumors from both early and late disease time points were significantly more viscoelastic than normal mammary glands. Spatial microindentation revealed that while local stiffness and viscoelasticity are strongly coupled in soft regions of normal tissue, this relationship weakens in late carcinoma. Compared to normal glands, carcinoma tissues also exhibited marked upregulation of glycosaminoglycans (GAGs) and glycoproteins. Selective enzymatic degradation of sulfated GAGs, mucins, or O/N-linked glycans reduced viscoelasticity without altering stiffness, establishing glycoconjugates as major contributors to tumor viscoelasticity. In vitro, increasing matrix viscoelasticity drove non-transformed mammary epithelial cells (MCF10A) toward a malignant phenotype through focal adhesion kinase mechanosignaling. Our work establishes increased viscoelasticity as a defining mechanical feature of breast cancer progression, driven in part by the accumulation of glycoconjugates. These findings provide a fuller picture of tumor mechanical evolution and a benchmark for recapitulating matrix viscoelasticity in engineered models of breast cancer.

## Introduction

During breast cancer progression, the tumor extracellular matrix (ECM) undergoes significant remodeling, resulting in changes to the mechanical properties of the tumor^1,2^. Increases in elastic modulus, or stiffness, of breast tumors are well recognized^3,4^, to the extent that manual palpation can be effectively used for screening. Breast tissue is not purely elastic but rather viscoelastic, exhibiting time-dependent mechanical responses such as viscous relaxation of stress in response to loading^5^. ECM viscoelasticity influences key cellular behaviors including differentiation^5,6^, morphogenesis^7,8^, proliferation^9,10^, and cancer cell invasion^11,12^. Increased ECM viscoelasticity, or faster stress relaxation, can promote aberrant cell shape^13,14^, dynamic protrusions^12^, and altered transcriptional programs^11,13,15^ in both breast cancer cells and non-transformed mammary epithelial cells. While it is clear that matrix viscoelasticity can influence breast cancer cell behavior, how viscoelasticity varies through breast cancer progression, and on what molecular basis, is currently unknown.

Shear rheology of bulk breast tumors has revealed a viscoelastic/viscoplastic mechanical profile^10,12,16^. Similarly, non-invasive imaging techniques such as shear wave and magnetic resonance elastography have reported increased tissue fluidity, indicating more liquid-like behavior in malignant breast masses compared to benign tissue^17–19^. However, whether tumor viscoelasticity is altered compared to normal mammary gland tissue, and how viscoelasticity evolves during breast cancer progression, are still open questions. Further, the spatial resolution of the prior measurements is on the order of millimeters, obscuring any local variation. Tumors are mechanically heterogeneous in this regime^20^, and cells respond to their immediate microenvironment rather than to bulk-averaged tissue properties. Thus, there is a critical need for viscoelastic characterization of breast tissue through different stages of progression and at multiple length scales.

Remodeling of the tumor microenvironment during breast cancer progression involves increased fibrillar collagen production, crosslinking, and fiber organization, resulting in elevated stiffness^1,17,21^. Our understanding of which ECM components might contribute to tumor viscoelasticity is limited. However, there is evidence for glycoconjugates such as proteoglycans and glycoproteins driving viscoelastic mechanical profiles in other tissues^22–25^. These classes of glycoconjugates are also upregulated in breast tumors^26–30^, providing an intriguing foundation for molecular remodeling that could drive viscoelastic changes. Furthermore, whether changes in stiffness and viscoelasticity are driven by the same molecular remodeling events, and in the same locations, or are independently regulated remains unknown.

Here, we utilized the MMTV-PyMT mouse breast cancer model to map viscoelastic changes at the bulk and microscale levels. We demonstrated that tissue viscoelasticity increased with disease progression and showed reduced association with tissue stiffness, with glycoconjugate accumulation acting as a key contributor to this mechanical shift. We further showed that tumor-mimicking viscoelastic changes alone drove malignant traits in cells in vitro. Together, these findings define viscoelasticity as a mechanical axis of tumor progression distinct from stiffness, and provide the benchmark needed to help design more accurate breast cancer models.

## Results

We used the MMTV-PyMT genetically engineered mouse model to recapitulate early and late stages of breast cancer progression observed in humans. In this model, female Friend virus B (FVB) mice expressing the PyMT antigen develop progressive mammary tumors as they age^31^. Tumor samples from 8-9 week-old mice (early carcinoma) and 10-12 week-old mice (late carcinoma) were excised for mechanical and histological characterization (Fig. 1a). As controls, normal mammary glands were extracted from 8-10 week-old mice that were not carriers for the transgene and thus did not develop tumors. Hematoxylin and eosin (H&E) staining showed filling of ductal lumens and increased cell density in early carcinoma, and invasion into the surrounding tissue in late carcinoma samples, recapitulating aspects of human breast cancer progression in vivo^31,32^ (Fig. 1b).

**Fig. 1:**
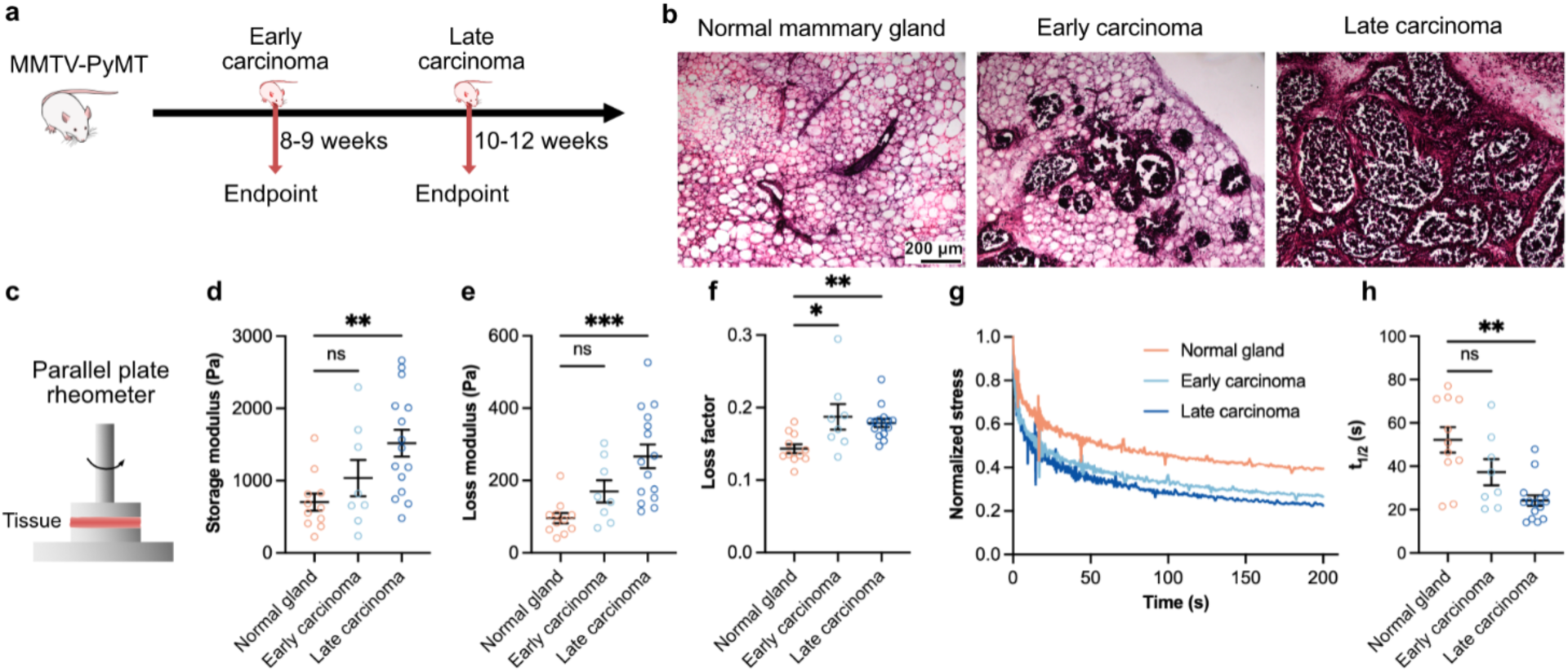
MMTV PyMT mouse mammary tumors display increased bulk viscoelasticity. **a**, Mice carrying the PyMT oncogene were allowed to develop tumors until weeks 8-9 (early carcinoma) and weeks 10-12 (late carcinoma). Non-carrier mice aged 8-10 weeks were used for normal glands. **b,** H&E histology showed normal mammary ducts in normal glands, increased epithelial cellularity and filling of ductal lumen in early carcinoma, and invasion into surrounding tissue in late carcinoma. **c,** Schematic of shear rheometry. **d,e,** Late carcinoma samples had a significantly higher storage modulus (**d**) and loss modulus (**e**) than normal glands, measured at 0.5% strain and 1 Hz. **f,** Both early and late carcinoma tissue also had a significantly higher loss factor (tan δ) than the normal glands. **g,** Representative stress relaxation profiles of 6 mm diameter samples tested at a 10% strain. **h,** Late carcinoma samples also showed a significantly shorter time to reach half of the maximum stress value (t1/2) than the normal glands. For **d–f,h,** n≥2 independent glands per mouse, with at least 3 mice per group; mean ± s.e.m.; Kruskal-Wallis test followed by post-hoc Dunn’s multiple comparison test was performed. ∗p < 0.05; ∗∗p < 0.01; ∗∗∗p < 0.001, not significant = ns.

To identify changes in bulk tissue mechanical properties during disease progression, we performed oscillatory shear rheology on tumor and normal tissue samples (Fig. 1c). Compared to normal glands, both early and late carcinoma samples had elevated storage and loss moduli, but this increase was significant only in the late carcinoma samples (Fig. 1d,e). The loss factor, defined as the ratio of loss modulus to storage modulus, which reflects the material’s ability for viscous dissipation, was significantly higher in both early and late carcinoma samples compared to the normal glands (Fig. 1f). Stress relaxation, another commonly used measure of material viscoelasticity, was assessed by applying a constant 10% strain to the samples and measuring the shear stress over time. The stress relaxation time (t_1/2_) was quantified as the time taken for the stress to reach half of its initial maximum value^5,33,34^. Both early and late carcinoma tumors relaxed stress faster than the normal glands, though only late carcinoma tumors were significantly faster (Fig. 1g,h). The increased loss factor and faster stress relaxation in tumors indicate that, at the bulk tissue level, mammary tissues exhibit increased viscoelasticity during disease progression.

Bulk rheological testing cannot resolve mechanical heterogeneity within a tumor, and it does not capture the mechanical properties of the local environment that the cells sense. To investigate changes in the local tissue mechanics at the microscale, we performed microindentations using a probe of diameter 55 μm and an indentation depth of 2 μm on normal glands, as well as early and late carcinoma tissues (Fig. 2a). The microindentation measurements showed that the Young’s modulus, obtained by fitting the Hertz contact model over the indentation loading curve (Fig. 2b), was significantly higher for both early and late carcinoma samples compared to the normal glands (Fig. 2c). Further, the loss factor, derived using dynamic mechanical analysis (DMA) during indentation at a frequency of 0.5 Hz and an amplitude of 200 nm, was significantly higher in both early and late carcinoma samples than in normal glands, indicating increased viscous dissipation (Fig. 2d). We also performed stress relaxation tests on the samples by holding the indentation depth constant for 200 s, which was chosen based on the range of stress relaxation times observed in bulk testing of the samples. Similar to our bulk rheological measurements, both early and late carcinoma samples exhibited faster stress relaxation profiles and faster relaxation times (t_1/2_) compared to the normal glands (Fig. 2e,f). Since relaxation exceeded the 200 s measurement window in many indentations, we quantified the percentage of indentations per sample with t_1/2_ > 200 s. The percentage of these indentations fell from 73% in normal glands to 50% in early and 20% in late carcinoma, reaching significance only for late carcinoma (Fig. 2g). As an alternative to stress relaxation time, we quantified the extent of stress dissipation after 200 s, which we report as the residual stress normalized to the maximum stress. Tissue that dissipates more stress would exhibit a lower residual stress at the end of the relaxation time window. Both early and late carcinoma samples displayed a significantly lower residual stress than normal glands, demonstrating increased viscoelasticity at the microscale during tumor progression (Fig. 2h). Overall, our microindentation findings reveal that early and late carcinoma tissues are more viscoelastic at the microscale, exhibiting faster stress relaxation and lower residual stress than normal mammary glands.

**Fig. 2:**
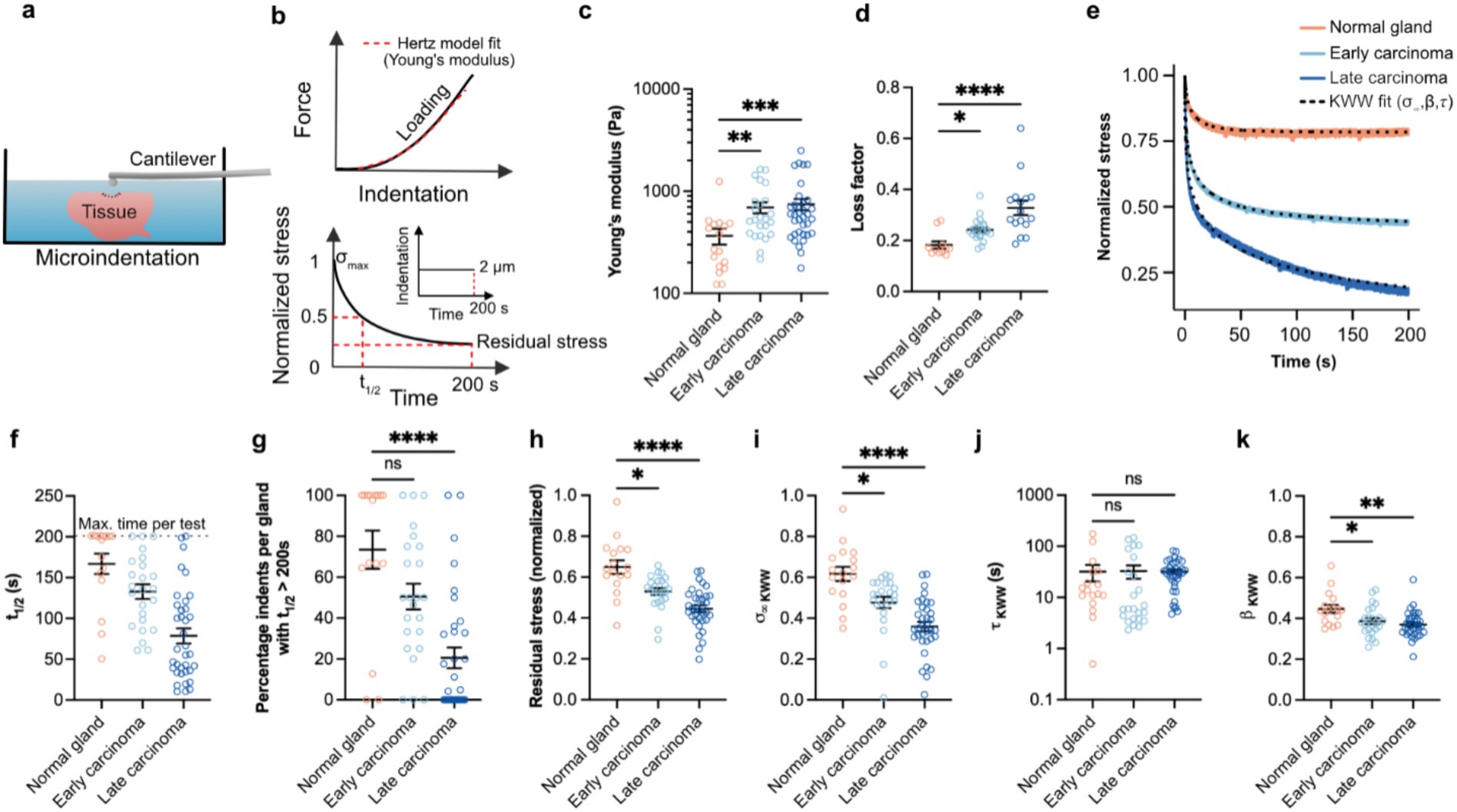
Tumor progression is associated with increased viscoelasticity at the microscale. **a**, Tissues immersed in PBS were tested using a microindenter with an indentation depth of 2 µm. **b,** Young’s modulus was determined from the loading curves using the Hertz contact model. Stress relaxation tests were performed by holding a constant indentation for a duration of 200 s. **c,** Both early and late carcinoma tumors showed a significantly higher Young’s modulus compared to the normal glands. **d,** Loss factor was significantly higher in early and late carcinoma tumors than normal glands. **e,** Representative stress relaxation profiles for samples and Kohlrausch-Williams-Watts (KWW) viscoelastic model fits. **f,** Time to reach half of the maximum stress value (t1/2), with maximum time values capped at 200 s. **g,** Late carcinoma showed a significantly lower proportion of indentations per sample that had a relaxation time (t1/2) > 200 s than normal glands. Samples with at least 4 indentations were considered. **h,** Both early and late carcinoma samples showed significantly reduced residual stress at 200 s, normalized to the peak stress. KWW viscoelastic model fit over the stress relaxation data showed that in comparison to the normal glands, **i,** σ∞ KWW, the equilibrium stress in early and late carcinoma samples was significantly lower, while **j,** *τ*KWW, the characteristic relaxation time showed no significant differences, and **k,** βKWW, the stretching exponent, was also significantly lower in early and late carcinoma samples. For **c, d, f, h-k,** n ≥ 3 independent tissue samples per mouse, with at least 3 mice per group, and at least 3 indentations per sample were performed. For **c, d, f-k,** mean ± s.e.m.; Kruskal-Wallis test followed by post-hoc Dunn’s multiple comparison test was performed. ∗p < 0.05; ∗∗p < 0.01; ∗∗∗p < 0.001; ∗∗∗∗p < 0.0001, not significant = ns.

To resolve which features of the stress relaxation response changed with disease progression, we fit the stress relaxation indentation data to the Kohlrausch-Williams-Watts (KWW) viscoelastic model. The KWW model is commonly applied to stress relaxation in complex heterogeneous materials such as tissues and hydrogels. It is governed by three parameters: equilibrium stress (σ_∞ KWW_), a characteristic relaxation time (*τ*_KWW_) defining the time for the relaxation function to decay to e^-1^ of the initial value, and a stretching exponent (β_KWW_) bound between 0 and 1 that reflects the range of contributing timescales driving relaxation^35–37^. We found that the equilibrium stress, representing the stress stored elastically in the tissue, was significantly lower in both early and late carcinoma groups compared to the normal glands (Fig. 2i). The characteristic relaxation time (*τ*_KWW_) showed no significant differences across the different groups (Fig. 2j), indicating a preserved timescale of relaxation during disease progression. In contrast, β_KWW_ was significantly lower in both early and late carcinoma samples compared to the normal glands (Fig. 2k). This decrease in β signifies a greater deviation from a standard linear solid (SLS), suggesting the presence of multiple stress relaxing elements in the matrix. Together, the KWW fit indicates that carcinoma tissues dissipate more stress and relax through a more stretched profile than normal glands, without a change in the characteristic relaxation time.

While spatial heterogeneity in stiffness is well established in malignant mammary tumors^4,38,39^, the spatial distribution of viscoelasticity is not well characterized. To map these spatial variations, we performed a scan of indentations across a 1 mm × 1 mm grid with 250 µm spacing between adjacent points, simultaneously measuring both Young’s modulus and the residual stress (at t = 200 s) (Fig. 3a). Representative heatmaps revealed spatial variation in both properties within each sample, alongside an overall increase in Young’s modulus and decrease in residual stress with tumor progression (Fig. 3b,c). Next, we investigated whether Young’s modulus spatially covaries with viscoelasticity, leveraging microindentation to overcome the confounding effects of spatial heterogeneity within the tissue. To study this relationship, we plotted residual stress against Young’s modulus for every indentation, using a moving average with a ±0.1 log-unit window to smooth the local trend (Fig. 3d). The normal, early, and late carcinoma groups separated vertically across the full stiffness range, demonstrating that regions of equal local stiffness dissipated more stress in carcinoma than in normal glands.

**Fig. 3:**
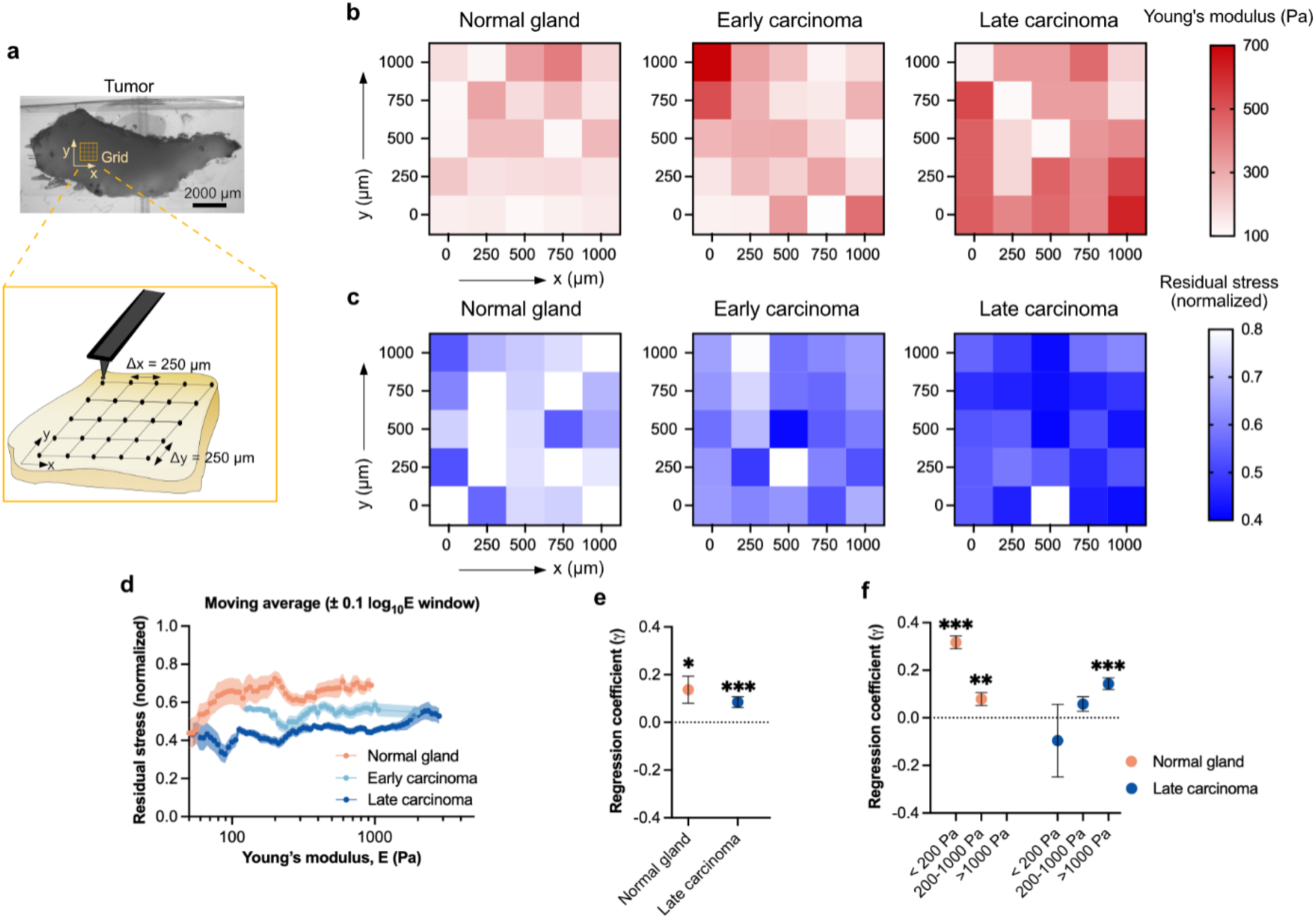
The relationship between stiffness and viscoelasticity is altered with disease progression. **a**, A raster-scan of indentations across a 1 mm × 1 mm grid with 250 µm spacing between adjacent points was performed on the samples. The Young’s modulus and residual stress (at 200 s) were measured at the same spatial coordinates. **b,** Compared to normal glands, carcinoma samples showed an increase in Young’s modulus and **c,** decrease in residual stress with disease progression. **d,** Moving average of residual stress plotted against Young’s modulus, using a ±0.1 log-unit window. The moving average profiles showed a clear distinction between groups along the residual stress axis. Glands with at least 3 indents per sample were considered; mean ± s.e.m. is plotted. **e,** Linear regression of residual stress on log-transformed Young’s modulus, with fixed effects for tissue sample, over the entire stiffness range showed a modest but significant relationship between the two variables. **f,** Normal glands showed a strong and significant association between stiffness and viscoelasticity in the < 200 Pa stiffness range. Late carcinoma samples showed only a weak association across the < 200 Pa, 200-1000 Pa, and > 1000 Pa ranges, with a significant regression only in the stiffest range of > 1000 Pa. For **e-f,** glands with at least 5 indents were considered; mean ± s.e.m.; Wald t-test with standard errors clustered by mouse. ∗p < 0.05; ∗∗p < 0.01; ∗∗∗p < 0.001, not significant = ns.

To quantify the association between stiffness and viscoelasticity, we performed a linear regression of residual stress on Young’s modulus. Regression analysis over the entire range of Young’s modulus showed that residual stress had a modest, though statistically significant, positive relationship with Young’s modulus in both normal glands (regression coefficient γ = 0.14) and late carcinoma samples (γ = 0.08) (Fig. 3e). The weak association is evident from the non-monotonic trend between the two variables in the moving average plots (Fig. 3d). We sought to further elucidate the relationship between stiffness and viscoelasticity by performing regression across three biologically relevant stiffness ranges: soft (< 200 Pa) corresponding to the healthy tissue; intermediate stiffness (200-1000 Pa) corresponding to the premalignant and tumor-adjacent tissue; and stiff (> 1000 Pa) corresponding to the malignant tissue^2–4^. In normal mammary glands, residual stress showed a strong, significant positive association (γ = 0.32) with stiffness in the soft range, indicating that higher local stiffness was associated with reduced viscoelasticity. A weaker but significant association (γ = 0.08) was seen in the intermediate stiffness regions within normal glands (Fig. 3f), and no indents exceeded 1000 Pa for the normal glands. However, late carcinoma samples showed only a weak association between stiffness and residual stress across the soft (γ = −0.09), intermediate (γ = 0.06), and stiff (γ = 0.14) ranges. The relationship was significant only in the stiff regions in the late carcinoma tissue. Together, our results show that stiffness and viscoelasticity are coupled differently in normal glands and carcinoma. In normal glands the two are tightly linked, with stiffer regions relaxing less, whereas in late carcinoma this coupling is weak. Regions that become stiffer in the tumor are therefore not consistently less viscoelastic.

Next, we sought to identify the molecular components within the tumors that contribute to changes in viscoelasticity during disease progression. Fibrillar collagen, proteoglycans, and glycoproteins are major constituents of the mammary ECM. Various proteoglycans and glycoproteins are known to be upregulated during tumor progression in the PyMT mouse breast cancer model^40,41^. Further, in humans, their upregulation is associated with poor patient outcomes in breast cancer^26,27,42^. High expression of fibrillar collagen is strongly associated with tumor progression in mice^1,3^ and with increased risk of breast cancer in humans^1,43^. We first tested whether proteoglycans and glycoproteins also vary in expression during disease progression in the mouse tumor model. Proteoglycans are molecules that contain a protein backbone, covalently attached to large and usually negatively charged glycosaminoglycans (GAGs). Using Alcian blue staining, we observed significant GAG accumulation in both early and late carcinoma tissues in comparison to the normal glands (Fig. 4a,b). We then examined specific GAGs and observed significantly higher levels of chondroitin sulfate, heparan sulfate, and hyaluronic acid in early and late carcinoma samples than in normal glands (Fig. 4c-e). The periodic acid–Schiff (PAS) stain, used to detect neutral carbohydrate-containing molecules such as glycoproteins, also showed a significantly greater intensity in both early and late carcinoma samples (Fig. 4a,f). Similarly, using picrosirius red staining for fibrillar collagen, we observed a significant increase in collagen fiber count, length, and width in late carcinoma samples compared to normal glands (Supp. Fig. 1a-d). Fiber coherency, a quantitative measure of local fiber alignment, ranging from 0 (isotropic) to 1 (highly aligned), was significantly higher in both early and late carcinoma samples than normal glands (Supp. Fig. 1e). These results highlight that tumor progression is associated with extensive ECM remodeling, involving upregulation of GAGs, glycoproteins, and fibrillar collagen.

**Fig. 4:**
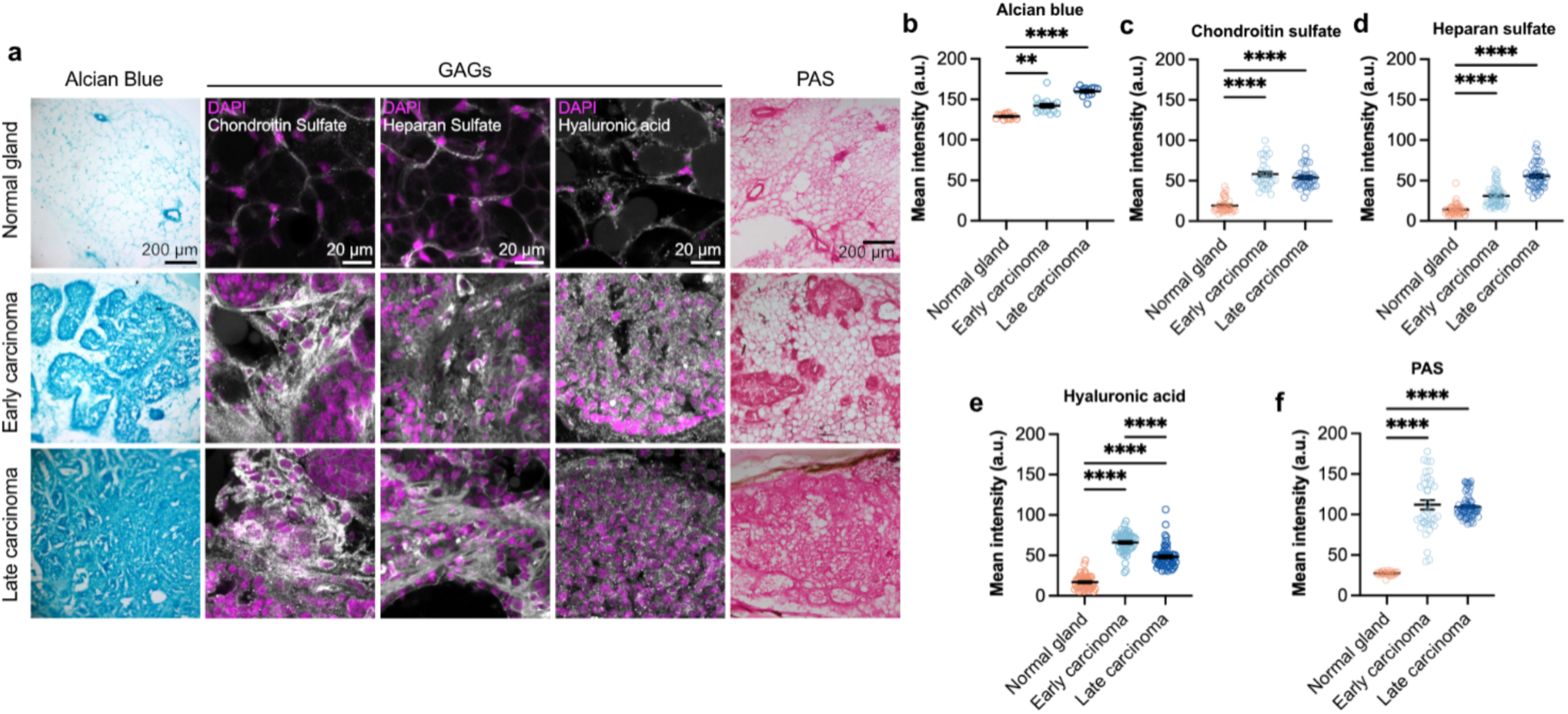
Tumor progression is associated with increased GAG and glycoprotein content in the tissue. **a**, Tissue sections were stained for GAGs using Alcian blue and using chondroitin sulfate, heparan sulfate, and hyaluronic acid antibodies. Glycoproteins were stained using the PAS stain. In comparison to normal glands, both early and late carcinoma samples showed a significantly increased mean staining intensity for **b,** Alcian blue, **c,** chondroitin sulfate, **d,** heparan sulfate, **e,** hyaluronic acid, and **f,** PAS. For **b,f,** n ≥ 5 images per tissue, with at least 3 independent tissue samples per group, and each tissue from a different mouse. For **c-e,** n ≥ 15 images per tissue, with at least 3 independent tissue samples per group, and each tissue from a different mouse. For **b-f,** mean ± s.e.m.; Kruskal-Wallis test followed by post-hoc Dunn’s multiple comparison test was performed. ∗∗p < 0.01; ∗∗∗∗p < 0.0001, not significant = ns.

While upregulation of fibrillar collagen in breast cancer is associated with increased stiffness in tumors^1,3,44^, the observed increase in GAGs and glycoproteins during tumor progression prompted us to ask whether these molecules might contribute to elevated viscoelasticity. To investigate the contribution of each GAG, we treated the late carcinoma tissue with specific GAG-degrading enzymes and measured stiffness and viscoelasticity before and after treatment. Chondroitinase ABC was used to digest chondroitin sulfate, heparinase I and III to digest heparan sulfate, while hyaluronidase type I-S was used to digest hyaluronic acid. An enzyme-free buffer served as the control. To minimize the effects of spatial heterogeneity, we indented the same spatial coordinates before and after 8 h of enzymatic digestion (Fig. 5a). Compared to the no enzyme control, degradation of either chondroitin sulfate or heparan sulfate led to a significant increase in residual stress, implying reduced tissue viscoelasticity (lower viscous dissipation) (Fig. 5b). Degradation of hyaluronic acid also caused an increased residual stress, but this change was not significant. Representative stress relaxation curves show the marked increase in residual stress at 200 s, indicating lower viscous dissipation, in enzyme-treated but not control samples (Fig. 5d–g). Interestingly, there were no significant differences in Young’s modulus upon degradation of any of the GAGs (Fig. 5c). To validate that the enzyme concentration and incubation time period led to effective GAG degradation, immunofluorescence was used to quantify GAG abundance. After degradation, we observed a significant decrease in signal intensity for chondroitin sulfate and hyaluronic acid; however, surprisingly, a significant increase in intensity was observed for heparan sulfate (Supp. Fig. 2a-d). It is possible that the elevated signal intensity results from increased accessibility of the epitope on the protein backbone of the proteoglycan to which the heparan sulfate antibody binds^45^. We also observed a reduction in Alcian blue staining after treatment with heparinase, confirming the degradation of heparan sulfate in the tissue section (Supp. Fig. 2e).

**Fig. 5:**
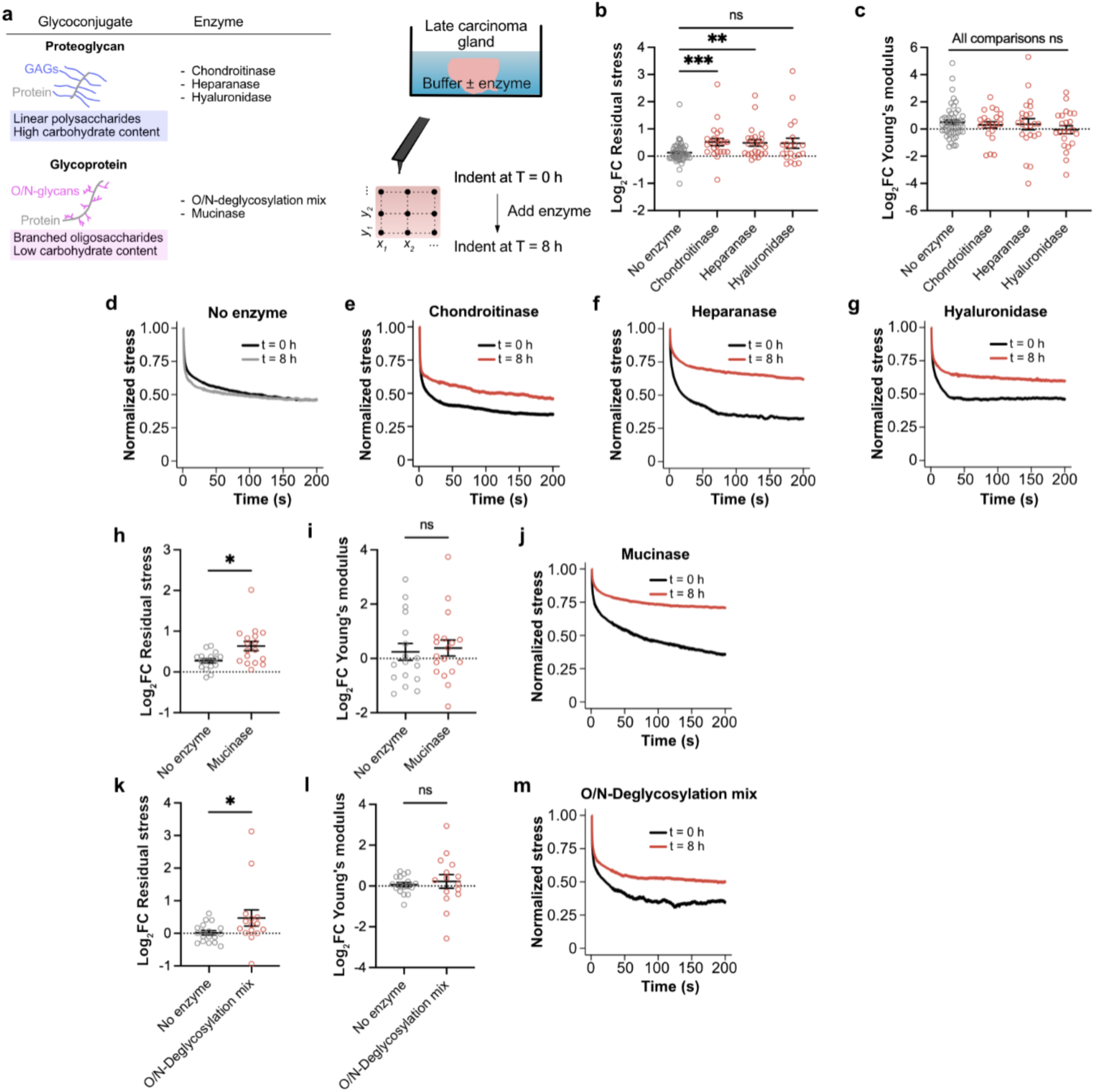
Enzymatic degradation of GAGs and glycoproteins reduces the viscoelasticity in late carcinoma tissue. **a**, Late carcinoma tissue samples were incubated with the GAG-degrading enzymes chondroitinase, heparinase, or hyaluronidase; or glycoprotein-degrading enzymes O/N-linked glycan deglycosylation mix, or mucinase for 8 h at 37°C. The same spatial location was indented before and after enzyme addition to the buffer to account for spatial heterogeneity. Samples in buffer solution without enzyme were used as controls. **b,** A significant increase in residual stress (at 200 s) was observed upon treatment with chondroitinase or heparinase, indicating reduced stress relaxation in the tissue. **c,** Treatment with GAG-degrading enzymes led to no significant change in tissue Young’s modulus. **d-g,** Representative stress relaxation profiles at the same spatial location before and after treatment with GAG degrading enzymes. **h,** Treatment with mucinase (StcE) or **k,** O/N-Deglycosylation mix also led to a significant increase in residual stress but no significant change in Young’s modulus **i l**. **j,** Representative stress relaxation plot before and after treatment with mucinase or **m,** O/N-Deglycosylation mix shows reduced viscous dissipation. For **b, c, h, i, k, l,** 5 indentations were performed per tumor, with at least 3 independent tumors used per group, and each tumor derived from a different mouse (mean ± s.e.m.; Kruskal-Wallis test followed by post-hoc Dunn’s multiple comparison test was performed). ∗p < 0.05; ∗∗p < 0.01; ∗∗∗p < 0.001, not significant = ns.

We next examined whether glycoproteins contribute to tumor viscoelasticity. To this end, we used a mucin-selective protease, StcE (secreted protease of C1 esterase inhibitor), to degrade the protein backbone of mucins^25,46,47^, a class of heavily glycosylated glycoproteins. With mucin degradation, we also saw a significant increase in residual stress compared to the no-enzyme control (Fig. 5h,j), but no significant change in tissue Young’s modulus (Fig. 5i). To investigate the contribution of glycans to tumor viscoelasticity, we used a commercially available blend of O-glycosidase, PNGase F, neuraminidase, β1-4 galactosidase S, and β-N-acetylhexosaminidase to perform O/N-linked deglycosylation of glycoproteins, similar to previous studies^48,49^. After treatment, we observed a significant increase in residual stress compared to the no-enzyme control (Fig. 5k), with no significant change in tissue Young’s modulus (Fig. 5l). The higher residual stress after treatment was also reflected in the representative stress relaxation profile (Fig. 5m). We confirmed the O/N-linked deglycosylation and mucin degradation by reduced PAS staining in enzyme-treated sections compared to the no-enzyme controls (Supp. Fig. 2f,g). Overall, our results show that both long polysaccharide chains such as GAGs and highly glycosylated glycoproteins are upregulated in tumor progression and play a role in driving increased viscoelasticity in the diseased tissue.

Given that fibrillar collagen promotes increased matrix stiffness during tumor progression, we examined whether it also dictates tissue viscoelasticity. Treatment of the late carcinoma samples with collagenase I for 4 h to degrade fibrillar collagen led to a significant decrease in tissue Young’s modulus (Supp. Fig. 3a). However, unlike GAG or glycoprotein degradation, we observed a significant decrease in residual stress upon collagen degradation, indicative of increased tissue viscoelasticity (Supp. Fig. 3b). We further validated these findings using a 2 mg/ml reconstituted collagen I gel and similarly observed a reduction in both Young’s modulus and the residual stress in gels after 1 h of collagenase I treatment (Supp. Fig. 3c,d). These findings indicate that while degrading fibrillar collagen increases carcinoma tissue viscoelasticity, degrading GAGs and glycoproteins reduces it.

Previous studies have illustrated that mammary epithelial cells sense and respond to changes in ECM stiffness through integrin clustering and focal adhesion formation^3,44,50^. Focal adhesion kinase (FAK) is a key downstream component of this process, becoming activated via autophosphorylation at Tyr397 following β1-integrin engagement, and has been implicated in malignant progression of the mammary gland^51–55^. Therefore, we asked whether the differences in both stiffness and stress relaxation rate we measured between normal and carcinoma tissue were reflected in altered β1-integrin and phosphorylated FAK (pFAK) abundance and localization in vivo. We observed an increase in β1-integrin abundance from normal glands to early carcinoma, and from early to late carcinoma tissue (Fig. 6a, Supp. Fig. 4). While pFAK levels were higher in normal glands (Supp. Fig. 4) than in early and late carcinoma tissues, there was a decrease in pFAK polarization (peripheral to interior) in late carcinoma compared to the early carcinoma samples (Fig. 6a). These findings demonstrate that changes in tissue stiffness and viscoelasticity during breast cancer progression are accompanied by increased β1-integrin abundance and a loss of pFAK polarization.

**Fig. 6:**
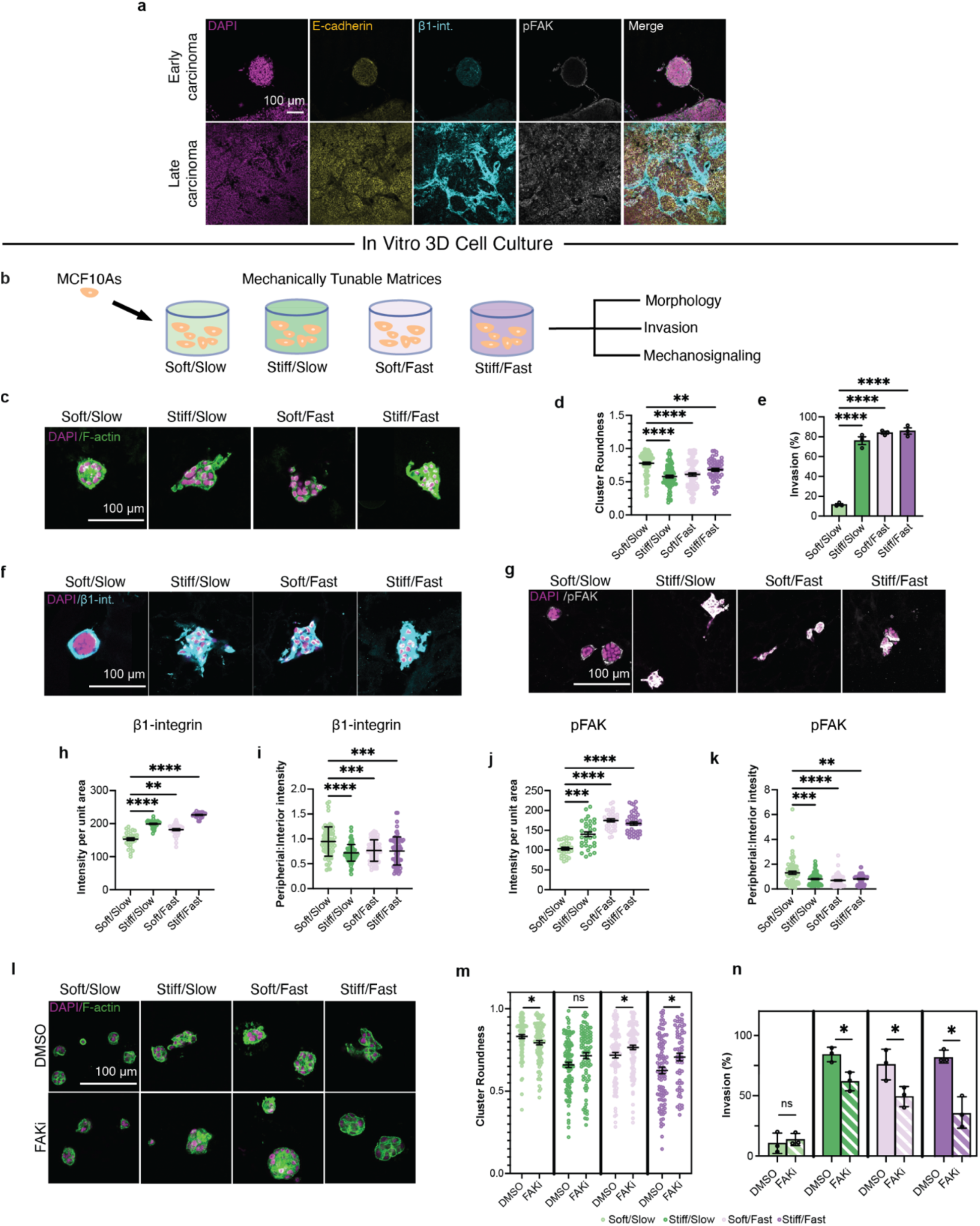
Mammary epithelial cells exhibit malignant traits in high viscoelasticity alginate-rBM matrices. **a**, Early and late carcinoma tissue stained for DAPI, E-cadherin, β1-integrin, and pFAK-Tyr397 showed a higher pFAK and β1-integrin expression in late carcinoma than early carcinoma tissue. **b,** MCF10A cells were encapsulated in alginate-rBM IPNs with tunable stiffness and viscoelasticity to investigate their morphology, invasion, and mechanosignaling. **c,** MCF10A cells were encapsulated in soft/slow-relaxing, stiff/slow-relaxing, soft/fast-relaxing, and stiff/fast-relaxing matrices and stained for DAPI and F-actin. **d,** Cluster roundness of cells was significantly lower in stiff and/or fast-relaxing conditions compared to the soft/slow-relaxing group. **e,** Percentage of invasive clusters was significantly higher in stiff and/or fast-relaxing conditions than in the soft/slow-relaxing group. **f, g,** Representative images of cells stained for DAPI and β1-integrin (f) or pFAK (g). **h,** Mean β1-integrin intensity was significantly higher in stiff and/or fast-relaxing conditions than the soft/slow-relaxing group. **i,** The peripheral:interior ratio of mean β1-integrin intensity was significantly lower in the stiff and/or fast-relaxing conditions than the soft/slow-relaxing group. **j,** Mean pFAK intensity, quantified from panel g, was significantly higher in stiff and/or fast-relaxing conditions when compared to the soft/slow-relaxing group. **k,** The peripheral:interior ratio of mean pFAK intensity was significantly lower in stiff and/or fast-relaxing conditions when compared to the soft/slow-relaxing group. **l,** MCF10A cells encapsulated across all mechanical conditions were treated with either DMSO (top) or FAK inhibitor (bottom), stained for DAPI and F-actin. **m,** Quantification of cluster roundness from panel l showed a significant increase upon treatment in both the soft/fast and stiff/fast groups. FAK inhibition significantly reduced roundness in the soft/slow group. **n,** The percentage of invasive clusters was significantly lower upon treatment across all mechanical groups except for soft/slow relaxing matrices. All scale bars are 100 μm. Statistical significance was tested using a Kruskal-Wallis test followed by Dunn’s multiple comparisons test in **d, h-k**; using a one-way ANOVA followed by Dunnett’s multiple comparisons test in **e**; and using Welch’s t-test in **m, n**. not significant = ns, ∗p < 0.05, ∗∗p < 0.01, ∗∗∗p < 0.001 and ∗∗∗∗p < 0.0001.

After characterizing the mechanics of normal, early carcinoma, and late carcinoma tissue, we asked whether increased matrix viscoelasticity, characterized by faster stress relaxation, could drive mammary epithelial invasion in vitro. We utilized alginate-reconstituted basement membrane (rBM) interpenetrating networks (IPNs), a system where a bioinert and ionically crosslinked alginate network is interpenetrated with rBM (Fig. 6b), enabling independent tuning of stiffness and viscoelasticity (Supp. Fig. 5a-c), while presenting basement membrane proteins necessary for integrin engagement and apicobasal polarity. When MCF10A mammary epithelial cells were encapsulated in soft/slow-relaxing alginate-rBM IPNs, mimicking the mechanics of normal mammary glands, we found that they formed round, acinar structures (Fig. 6c,d). Cells encapsulated in matrices that mimic either the stiffness or stress relaxation rate of tumor tissues, or both (i.e., stiff/slow-relaxing, soft/fast-relaxing, or stiff/fast-relaxing), developed into aberrant, invasive structures, marked by a significant decrease in roundness relative to their soft/slow-relaxing counterparts (Fig. 6c,d). Additionally, we found a significantly higher percentage of invasive clusters in stiff and/or fast-relaxing matrices compared to the soft/slow-relaxing group (Fig. 6e).

Given that both stiffness and stress relaxation influenced epithelial cluster morphology and invasion, we next sought to identify the mechanotransduction pathway underlying this response. Since β1-integrin engagement and FAK activation are central to how cells sense and respond to matrix mechanics^3,56^, we assessed β1-integrin and FAK phosphorylated at Tyr397. We found that both pFAK and β1-integrin abundance increased significantly in stiff and/or fast-relaxing matrices compared to soft/slow-relaxing matrices (Fig. 6f–h,j). Additionally, we found that while both pFAK and β1-integrin localized to the periphery of acinar clusters in soft/slow-relaxing matrices, the tumor-mimicking matrices (stiff/slow-relaxing, soft/fast-relaxing, and stiff/fast-relaxing) had significantly lower peripheral:interior abundance ratios, indicative of a loss in their polarization within the cell clusters (Fig. 6f-k). This overall significant increase in abundance but decrease in peripheral:interior localization of both pFAK and β1-integrin indicates that pFAK and β1-integrin are being engaged throughout the cell clusters in stiff and/or fast relaxing groups, rather than only at the periphery of the cell clusters as seen in the soft/slow-relaxing condition. These data suggest that the broader activity of pFAK and β1-integrin is associated with the malignant behavior seen in the stiff and high viscoelasticity conditions.

Next, we inhibited FAK in this system using the small-molecule inhibitor PF-573228 to understand the role of focal adhesion signaling in regulating the malignant behavior in tumor-mimicking mechanical conditions. FAK inhibition led to a significant increase in roundness in soft/fast relaxing and stiff/fast relaxing matrices compared to DMSO vehicle controls (Fig. 6l,m). Interestingly, there was a small but significant decrease in roundness in the soft/slow relaxing matrices upon FAK inhibition, potentially pointing to a role for FAK in maintaining acinar integrity. Additionally, there was a significant decrease in the percentage of invasive clusters upon FAK inhibition compared to DMSO controls in all stiff and/or fast relaxing tumor-mimicking groups (Fig. 6n). Taken together, these findings show that an increase in matrix stiffness or viscoelasticity can independently drive malignant behavior in mammary epithelial cells. Further, this invasive response in mammary epithelial cells is regulated by FAK signaling in response to high tumor-mimicking viscoelasticity.

## Discussion

Using the MMTV-PyMT mouse breast cancer model, we show that both early and late carcinomas exhibit increased viscoelasticity compared to normal mammary glands, at both the bulk and microscale. Our findings are consistent with previous imaging-based studies that found that malignant lesions are more liquid-like compared to benign breast masses^17,18^, and may therefore undergo greater viscous dissipation. Another study reported higher fluidity in malignant versus benign human breast lesions using a Kelvin-Voigt fractional derivative model applied to atomic force microscopy (AFM) data^57^. This study probed timescales under 1 s, too short to resolve slower stress relaxation events such as ECM network rearrangement. Our stress relaxation measurements, conducted over longer timescales of up to 200 s, captured the range of stress relaxation times for common natural ECM polymers^58^.

While recent work has emphasized the importance of ECM viscoelasticity, the molecular components that contribute to regulation of tissue viscoelasticity remain underexplored. Using selective enzymatic degradation, we identified glycoconjugates, including GAG-containing proteoglycans and glycoproteins, as previously unrecognized contributors to tumor viscoelasticity. While the precise mechanism by which glycoconjugates such as proteoglycans mediate tissue viscoelasticity remains unclear, GAGs contribute to viscoelasticity in multiple tissue types^22,23,59,60^. Heavily glycosylated mucins also provide lubrication in ocular and cartilage tissues^25,47^. The negative charges on sulfated GAG chains retain water and can generate osmotic pressure through swelling, contributing to ECM viscoelasticity^24^. Apart from modulating viscous properties of the ECM, GAGs can also influence collagen fiber alignment, ECM tensile strength, and long-range force transmission^61,62^. Perlecan, a heparan sulfate proteoglycan, modifies the lifetime of collagen IV molecules in the basement membrane, and molecular dynamics simulations link collagen IV lifetime to the stress relaxation time of the network^63^. Further, shRNA-based knockdown of versican, a type of chondroitin sulfate proteoglycan, prevented the increase in viscoelasticity observed in pleural fibrosis^64^. At the cell surface, glycocalyx-associated glycoproteins were also upregulated in tumors, where bulky glycoproteins and mucins can alter plasma membrane mechanical properties, cell fluidity, and downstream integrin-mediated mechanotransduction^65,66^. The extent to which glycocalyx remodeling also contributes to tissue viscoelasticity remains to be explored, and will require cell-type-specific glycan perturbation studies.

Increased crosslinking of fibrillar collagen in the ECM is associated with elevated tumor stiffness in multiple cancer types^67^, but its relation to tissue viscoelasticity is less well characterized. In reconstituted collagen gels, crosslinking of collagen leads to slower stress relaxation and thus reduces viscoelasticity^58,68^. The positive association between stiffness and residual stress that we observed in stiff regions (> 1 kPa) of late carcinoma may therefore reflect increased collagen crosslinking. Conversely, we found that degrading fibrillar collagen increased viscoelasticity in both late carcinoma tissue and collagen gels. Both results indicate that crosslinking of the fibrillar collagen network reduces viscoelasticity while degradation of the network increases it. Matrix metalloproteinases degrade fibrillar collagen to promote metastasis in invasive carcinoma^69–71^, which may similarly lead to increased ECM viscoelasticity in the tumor microenvironment. Future work should examine how counterbalancing factors, such as collagen crosslinking versus enzymatic degradation of collagen, and glycoconjugate production, regulate tumor viscoelasticity over time.

Beyond their contribution to tissue mechanical properties, glycoconjugates are also upregulated in tumors and are associated with poor patient outcomes^26,28–30,72^. We found significantly higher levels of both GAGs and glycoproteins in early and late carcinoma samples compared with normal mouse mammary glands, consistent with reports that GAG-bearing proteoglycans are markedly upregulated in the tumor ECM of mouse breast cancer models^40,41,73,74^. High levels of proteoglycans such as syndecan and versican are linked to the malignant phenotype and poor prognosis in multiple cancer types^28,42,72^. High expression of the chondroitin sulfate proteoglycan CSPG4 is likewise associated with lower overall survival and shorter time to recurrence in breast cancer patients^26,75^. Increased hyaluronic acid levels are similarly considered a prognostic marker of disease progression and correlate with poor patient outcomes^29,76,77^. Among glycoproteins, overexpression of many cell surface and extracellular mucin subtypes is also linked to poor prognosis in breast cancer^27,30^. Furthermore, reducing the expression of proteoglycans, mucin 1, and aberrant protein glycosylation can prevent breast cancer cell invasion^41,78–80^, establishing their functional role in driving malignant traits. It remains to be studied how increased density of glycoconjugates and the resulting increased tissue viscoelasticity may synergize in driving disease progression.

We showed that enzymatic degradation of GAGs, O/N-linked glycans, and mucins regulates tissue viscoelasticity largely independently of stiffness. Stiffness and viscoelasticity therefore have distinct molecular determinants in the late carcinoma tissue. Further, the association between stiffness and viscoelasticity is strong in normal glands but weakens in late carcinoma, implying that stiffness does not reliably predict viscoelasticity in tumor tissue. Cells in different tumor regions may therefore encounter mechanical states that stiffness measurements alone would not reveal.

Whether stiffness and viscoelasticity act through shared or distinct mechanotransduction pathways in cancer cells remains an open question. It is well established that increased matrix stiffness can drive a malignant phenotype in vitro^3,21,44,81–85^. Studies have also explored how viscoelasticity can influence cell behavior in 3D in vitro systems^7,11,13,33,86–88^, and a smaller number have specifically shown that altered viscoelasticity impacts mammary epithelial phenotype and invasion^7,11,13,89^. In agreement with previous literature^5,7,8,11,50,56,81,86,90^, we found that both increased stiffness and altered stress relaxation rate disrupted acinar morphology, promoted invasive cluster formation, and increased β1-integrin expression, and that the combination of the two produced the highest percentage of invasive clusters and β1-integrin expression. The increased abundance of β1-integrin and loss of pFAK polarization in fast-relaxing matrices were also observed in the late carcinoma tumors.

While the role of FAK has been well established in driving invasive phenotypes of mammary epithelial cells in stiff matrices^50,91^, there is limited knowledge of FAK mechanosignaling in stress relaxing matrices^7,8,92^. Here, we saw that FAK inhibition reduced the malignant phenotypes seen in matrices with faster stress relaxation. Fast stress relaxing alginate networks can be physically remodeled^5^ and thus may enable integrin clustering, focal adhesion formation, and FAK signaling. Future mechanistic studies should investigate whether stiffness and viscoelasticity drive mammary epithelial invasion through shared or distinct signaling pathways.

In summary, using a mouse model of breast cancer, we showed that tumor progression is associated with increased viscoelasticity, both in bulk and at the microscale. Since stiffness and viscoelasticity were measured at the same location, we could demonstrate that the association between these two mechanical properties was weaker in the carcinoma tissue compared to the normal mammary gland. Mechanistically, this work also revealed that the upregulation of glycoconjugate molecules such as GAG-bearing proteoglycans and glycoproteins during disease progression contributed to increased tumor viscoelasticity, independently of stiffness. Further, using hydrogel models with tunable viscoelasticity, we showed that faster stress relaxation of the ECM can lead to direct changes in β1-integrin and pFAK-based mechanotransduction signaling pathways. While validation in human tissue is still required, our findings identify viscoelasticity as a distinct and independently varying mechanical property of tumor progression. Given that glycoconjugate upregulation and ECM remodeling occur across solid tumors^93^, changes in tissue viscoelasticity likely accompany progression in other cancers. Thus, our findings motivate broader viscoelastic characterization of solid tumors and incorporation of viscoelasticity into in vitro disease models for drug screening and therapeutic design.

## Methods

### Animal studies

All animal experiments were conducted according to the experimental procedures approved by the Institutional Animal Care and Use Committee at University of California, Santa Barbara. Hemizygous or non-carrier female FVB/N-Tg(MMTV-PyVT)634Mul/J mice were purchased from the Jackson Laboratory (Cat # 002374).

### Tissue Rheology

Fresh tissue from mice was harvested and tested on an Anton Paar MCR-502e strain-controlled rheometer. Fresh samples were prepared using 6 mm diameter biopsy punches and stored in PBS until testing. Storage modulus, loss modulus, and loss factor (tan δ) were measured at 0.5% strain and at a frequency of 1 Hz. Stress relaxation tests were performed at a 10% strain and the time required for the stress to reach 50% of the maximum value was recorded.

### Microindentation

Fresh tissue samples were maintained in PBS supplemented with 1% penicillin/streptomycin and protease inhibitor cocktail without EDTA (GenDEPOT, Cat #P3100). Samples were embedded in 4% w/v agarose and cut using a microtome blade to expose a flat surface for indentation. These samples were adhered to the base of a 6-well plate using superglue and washed multiple times with 1X PBS to remove any debris. Samples were tested on the same day for microindentation using an Optics11 Pavone microindenter at the BioPACIFIC MIP user facilities at UC Santa Barbara. Measurements were performed using a cantilever with a stiffness of 0.47 N/m and a spherical probe of radius 27.5 µm, chosen for the intended range of tissue stiffness according to the instrument specifications. The sample was loaded and unloaded at a speed of 10 μm/s to keep the indentation time much smaller than the material’s relaxation time. Young’s modulus was calculated by using a target indentation of 2 µm and fitting the Hertz contact model to the loading curve using the equation:

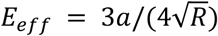

where *a* = *F*/*δ*^3/2^ ; *R* = tip radius, *F* = load, δ = indentation

Young’s modulus, *E_sample_* is obtained from *E_eff_* using,

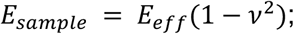

assuming Poisson’s ratio *ν* = 0.5.

Upon reaching the final indentation depth, the indentation depth was held constant in indentation-control mode for a dwell time of 200 s to perform the stress relaxation test. To measure the loss factor, dynamic mechanical analysis (DMA) was performed at 0.5 Hz at an oscillation amplitude of 200 nm superimposed on the held indentation depth, spanning approximately 10 oscillation cycles per measurement window. The load and indentation signals were independently fit in R using nonlinear least squares (nls) to sinusoidal functions:

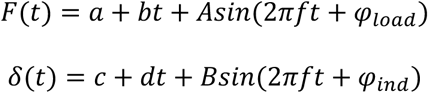

where *a* and *c* are offsets, *b* and *d* are linear drift terms included to account for viscoelastic creep during oscillation, *A* and *B* are the fitted oscillation amplitudes, *f* is the applied frequency, and *φ_load_* and *φ_ind_* are the phase angles of the load and indentation signals respectively. The loss factor was calculated as:

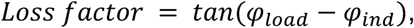

where (*φ_load_* − *φ_ind_*) is the phase lag.

The Kohlrausch-Williams-Watts (KWW) viscoelastic model was fit to the tissue stress relaxation data by nonlinear least squares using the equation:

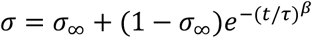

where σ = stress normalized to the peak stress, *σ*_∞_ = normalized equilibrium stress, β = stretching exponent bound between 0 and 1, and *τ* is the characteristic viscous relaxation time (s). For fitting, bounds of 0.1 to 200 s; 0 to 1; and 0 to 1 were chosen for *τ*, β, and *σ*_/_ respectively. A lower β indicates a broader distribution of stress relaxation times, while β = 1 represents a standard linear solid model.

### Moving average visualization of residual stress with respect to Young’s modulus

We calculated a sliding window moving average of normalized residual stress at 200 s along a log_10_-transformed Young’s modulus (E_sample_). Samples with at least 3 indentations were considered. Analysis was performed using a sliding window of ±0.1 log-units and a step size of 0.025 log-units. To prevent highly indented samples from disproportionately weighting local trends, individual indentation measurements were first averaged within each sample in a given window. Mean and standard error were then calculated within that window. Windows failing to meet a minimum threshold of 10 total indentations or two distinct tissue samples were omitted from analysis.

### Linear fixed-effects regression model of residual stress on Young’s modulus

Young’s modulus values were first log_10_-transformed. We then performed linear regression of normalized residual stress at 200 s on log_10_-transformed Young’s modulus (E_sample_) across distinct stiffness regimes (< 200 Pa, 200-1000 Pa, and > 1000 Pa). We restricted our analysis to samples that had at least 5 indentations recorded per sample. Models were fit separately for normal glands and late carcinoma using standard linear models (lm function in R). To isolate localized mechanical relationships from sample variation, we included fixed effects for individual tissue samples. To account for statistical clustering from multiple measurements per mouse, cluster-robust standard errors and Wald *t*-tests were calculated at the mouse level using the sandwich and lmtest packages in R.

### Mammary Epithelial Cell Culture

MCF10A mammary epithelial cells were obtained from American Type Culture Collection (ATCC) and cultured according to established protocols^7,94^. Prior to encapsulation, cells were grown as a monolayer in standard 2D tissue culture flasks. An established growth medium recipe was used: DMEM/F12 (Thermo Fisher Scientific, 11320033) basal medium was supplemented with 5% horse serum (Thermo Fisher Scientific, 16050122), 1% penicillin/streptomycin (Thermo Fisher Scientific, 15140122), epidermal growth factor (20 ng/ml; Peprotech, AF-100-15-1MG), hydrocortisone (0.5 μg/ml; Sigma-Aldrich, H0396), cholera toxin (100 ng/ml; Sigma-Aldrich, C8052), and insulin (10 μg/ml; Sigma-Aldrich, I1882). MCF10A cells were harvested for encapsulation by first washing with PBS and trypsinizing with 0.05% Trypsin/EDTA (Thermo Fisher Scientific, 25300054). Cells were pelleted at 200 rcf for 5 min, counted, and resuspended in DMEM/F12 to create a single-cell suspension. The suspension was added to the hydrogel precursors to establish a final seeding density of 5 × 10^4^ cells/ml.

### Alginate-rBM IPN fabrication and encapsulation of MCF10As

Dissolved alginate (0.5% or 1% w/v final concentration for LF20/40 or VLVG, respectively) in DMEM/F12, Matrigel (Corning, 354230) at a final concentration of 4 mg/ml, and a single-cell suspension (final concentration of 5 × 10^4^ cells/ml) in growth media were mixed in a Luer lock syringe. To initiate cross-linking of the alginate matrices, a calcium sulfate slurry (CaSO_4_ dissolved in sterile MilliQ water) was mixed with DMEM/F12, which was then introduced into a second, accompanying Luer lock syringe. Final calcium concentrations for soft and stiff matrices were 2 and 21 mM for LF20/40 (at 0.5 w/v%) matrices and 5 and 25 mM for VLVG (at 1 w/v%) matrices, respectively. The two syringes were coupled using a female-to-female Luer-lock coupler, and the contents were homogenized by passing them back and forth several times. The mixed hydrogel solution was rapidly cast into a 48-well plate at a volume of ∼500 μl per well. Following a 2-hour incubation at 37°C to allow complete cross-linking, growth medium was added, and the cultures were maintained for 7 days with media replacements every 1 to 2 days.

### Rheological testing of alginate-rBM IPNs

Gels were prepared as described above and cast directly between 25-mm-diameter parallel plates on an Anton Paar MCR-502e strain-controlled rheometer. The edges of the gel were sealed using mineral oil to prevent solvent evaporation. The gels were allowed to polymerize and their storage modulus was recorded every minute at 0.5% strain and 1 Hz frequency until the equilibrium modulus was reached. Upon reaching a constant equilibrium storage modulus, the stress relaxation test was then performed at a 10% strain. The time needed for the stress to reach half of the maximum value was then recorded. All the tests were performed at 25°C.

### FAK inhibition

The FAK inhibitor, PF-573228 (Cayman Chemical, 14924), was dissolved in DMEM/F12 with a final concentration of 5 μM. The same volume of DMSO was diluted in DMEM/F12 for the vehicle control. Inhibitor or DMSO solutions were added to growth media on day 1 of culture for alginate-rBM IPNs.

### Tissue and IPN histology

Tissues and IPNs were fixed in 4% formaldehyde in PBS for 1 h at 37°C. Tissues were then washed twice with PBS and incubated in a 30% w/v solution of sucrose in PBS overnight at 4°C. Cryoprotected samples were then incubated in a 1:1 solution of OCT and 30% w/v sucrose solution in PBS for 4-6 h at 4°C. Samples were then frozen in OCT in cryomolds. Frozen tissues were cut into 25-30 µm slices while frozen IPN samples were cut into 40 µm slices using a Leica Cryostat CM1850. Sections were placed on poly-L-lysine coated glass slides. Sections were first washed with PBS to remove OCT.

For Alcian blue staining, sections were rehydrated with 3% acetic acid solution for 3 min. Then samples were stained using a 1% w/v Alcian blue solution (ThermoFisher Scientific, Cat # J60122.09) in 3% acetic acid (pH 2.5) for 45 min at room temperature. The sections were rinsed with acetic acid twice for 1 minute per wash, followed by rinsing in tap water for 2 min. The samples were dehydrated first using 70% ethanol and then 100% ethanol for 2 min per rinse. The samples were then air dried and mounted using Permount (Fisher, Cat #SP15-100).

For Picrosirius red staining, samples were washed 2X with DPBS, 5 min per wash. Samples were then stained with 0.1% sirius red stain in picric acid (Electron Microscopy Sciences, Cat #26357-02) for 45 min at room temperature. Samples were then washed with 0.01 M HCl 4-5 times. Samples were then dehydrated sequentially using 70% and 100% ethanol, allowed to dry completely and then mounted using Permount.

For PAS staining, samples were rinsed in MilliQ water and then incubated with 0.55% periodic acid (Epredia, Cat #88016) for 8 min at room temperature. Samples were then rinsed in MilliQ water twice for 2 min and then incubated with the Schiff’s reagent (Epredia, Cat #88017) for 15 min at room temperature. Samples were then thoroughly rinsed under running tap water, then dehydrated sequentially using 70% and 100% ethanol. Samples were then allowed to air dry completely and then mounted using Permount. For samples stained with both Alcian blue and PAS, samples were stained with Alcian blue as described above, then rinsed with MilliQ water and then stained using the PAS protocol.

For H&E staining, rehydrated sections were stained with Hematoxylin 1 (Epredia, Cat #88018) for 1 min at room temperature and then rinsed twice with MilliQ water for 2 min per wash. Samples were then stained with 1% Eosin Y solution (Fisher, Cat # SE23-500D) for 1 min.

Samples were then rinsed quickly with MilliQ water for 1 min. Samples were then sequentially dehydrated with 70% and 100% ethanol, air dried, and then mounted with Permount.

### Immunofluorescence

PAP pen (Vector Laboratories, Cat #H-4000) was used to outline the gel or tissue section to create a hydrophobic barrier for antibody incubation steps. First, fixed tissue or gel sections were washed with PBS (+Ca^2+^) twice to remove OCT. Sections were blocked and permeabilized for 1 h at room temperature using a solution of 1× DPBS (with Ca^2+^/Mg^2+^), 1% bovine serum albumin, 0.1% Triton X-100, 0.3 M glycine, 10% goat serum, and 0.05% sodium azide. Sections were then incubated with primary antibody solutions overnight at 4°C. Sections were then washed 2X with the blocking buffer and incubated in Alexa Fluor 647 secondary antibody (1:1000), DAPI (1 μg/ml), and Alexa Fluor Phalloidin 488 (1:200, ThermoFisher Scientific) for 1 h at room temperature. Sections were then washed 3X using the blocking buffer, mounted using Prolong Gold antifade reagent (ThermoFisher Scientific), and the coverslip edges were sealed using transparent nail polish. Antibodies used were anti-chondroitin sulfate (1:25 dilution, DSHB, Cat #9BA12), anti-heparan sulfate (1:50 dilution, DSHB, Cat #33-2), anti-hyaluronic acid (1:100 dilution, Biomatik, Cat #CAU28389), β1 integrin (1:200, Thermo Fisher Scientific, 14-0299-82), FAK-P-Y397 (1:500, Thermo Fisher Scientific, Cat #700255), E-cadherin (1:500, BD Biosciences, Cat #610181), Alexa Fluor 488 goat anti-rabbit IgG, Alexa Fluor 555 goat anti-mouse IgG, Alexa Fluor 647 goat anti-rabbit, Alexa Fluor 647 goat anti-Mouse IgG2a, Alexa Fluor 647 goat anti-mouse IgM, and Alexa Fluor 647 goat anti-mouse IgG.

### Degradation of GAGs, Mucin, Collagen I, and O/N-linked Glycans

For all enzyme-based digestion assays, freshly harvested tissues were frozen in OCT using isopentane cooled over liquid nitrogen and thawed later. After thawing, samples were prepared for indentation as previously described. Samples were kept at 37°C in 1X HBSS buffer solution supplemented with 10 mM HEPES, 1% penicillin/streptomycin, and 1X protease inhibitor without EDTA during the entire experiment. Chondroitinase ABC (Cat #C2905), heparinase (Cat #H3917), hyaluronidase (Cat #H3506), and mucinase StcE (Cat #SAE0202) were used at a final concentration of 0.1 U/ml, 0.5 U/ml, 1.3 mg/ml, and 0.01 mg/ml respectively. All samples were incubated in buffer for 2 h followed by indentation. Buffer was then replaced by enzyme mix, except for control wells, and samples were incubated for 8 h and then indented again at the same x,y coordinates.

For O- and N-linked glycan removal, after thawing and embedding in agarose, samples were maintained in a 500 mM sodium phosphate buffer containing 1% penicillin/streptomycin and 1X protease inhibitor cocktail without EDTA at 37°C and pH 7.5 as recommended by the enzyme manufacturer. Protein Deglycosylation Mix II (New England Biolabs, Cat #P6044S) was used at 10 μl enzyme per 200 μl solution. Samples were incubated with the deglycosylation mix for 8 h at 37°C on the microindenter.

For collagen I digestion, after thawing and agarose embedding, tissue samples were incubated with PBS (containing Ca^2+^) and 1% penicillin/streptomycin at 30°C. 0.5 mg/ml collagenase (Sigma, Cat #C0130) was then added to the treatment wells and incubated for 4 h.

Collagen I gels were prepared using a 4 mg/ml collagen I stock solution (Advanced BioMatrix, Cat #5056) and diluted to 2 mg/ml in PBS (containing Ca^2+^) and neutralized using 1 M NaOH in a 12-well plate. Gels were allowed to form at 37°C for 1 h. Gels were incubated with either PBS or PBS containing 0.5 mg/ml collagenase I (Sigma, Cat #C0130) for 1 h at 30°C. Gels were then washed 2X with PBS and microindentation was performed on them.

### Image analysis

ImageJ (NIH) was used for image analysis. For analysis of signal intensity of Alcian blue and PAS stains, the integrated intensity computed for each image was obtained by converting each image to grayscale followed by thresholding above a fixed value across all images. The integrated signal density was then normalized to the thresholded area to obtain the mean signal intensity. At least 5 images per sample were acquired, and at least 3 independent samples from different mice, each for normal gland, early carcinoma, and late carcinoma groups were imaged.

For immunofluorescence images of chondroitin sulfate, heparan sulfate, and hyaluronic acid, at least 15 images per sample were acquired, and at least 3 independent samples, each from a different mouse were imaged. Mean fluorescence intensity was computed by measuring the integrated signal intensity using a fixed signal threshold across all images in ImageJ. This integrated signal intensity was then divided by the area above the threshold to obtain the mean signal intensity.

For measuring the mean intensity of the β1 and pFAK proteins in in vitro samples, the images were first thresholded to highlight the stain of interest. Next, fluorescence intensity was calculated by measuring the mean intensity per unit area (μm^2^) for each cluster. For measuring the peripheral to inner ratio of the mean signal intensity for β1 integrin and pFAK proteins in in vitro samples, we segmented each cluster from the β1 integrin channel and divided it into outer (peripheral) and inner (central) regions using morphological erosion/dilation in R (EBImage). Briefly, the cell mask was thresholded and hole-filled from the β1 integrin channel, and eroded to obtain a perimeter contour, which was then dilated to define a fixed-width outer band. The remaining central area was designated the inner region. Mean intensity within each region was calculated by applying the binary outer or inner mask to the corresponding fluorescence channel and averaging pixel intensity values within the masked area separately for the pFAK and β1 integrin channels.

### Collagen fiber dimensions and fiber alignment analysis

Picrosirius red stained samples were imaged using a 10X objective under polarized light on an Olympus BX60 upright compound microscope. Polarized light images of picrosirius red stained slides were processed through Fiji (ImageJ) to isolate the red channel only. Red channel images were then analyzed via CT-FIRE, a freely available software written in MATLAB script^95^, with the threshold set to 40, to quantify collagen fiber width, length, and number. At least 4 images per sample, and at least 3 independent samples of normal gland, early carcinoma, and late carcinoma were analyzed. The quantification for each image was averaged and reported as one data point. Collagen fiber coherency was measured via the OrientationJ macro^96^ in Fiji on the same red channel images described above. OrientationJ measures the local orientation of every pixel in an image, calculating a structure tensor by evaluating image intensity gradients. The structure tensor is summed over all pixels in the ROI and decomposed into its eigenvalues *λ*₁ and *λ*₂, where *λ*₁ ≥ *λ*₂. Coherency is defined as (*λ*₁ − *λ*₂) / (*λ*₁ + *λ*₂) and ranges from 0 to 1, where 0 indicates no preferred fiber alignment and 1 indicates perfect fiber alignment. Using the OrientationJ measure function, 3 ROIs were drawn and measured per image. ROIs were selected based on areas of high alignment by eye. For images with no apparent alignment, an ROI was drawn within a representative region. The coherency measurements from each image were then averaged and reported as one data point.

### Statistical analysis

Statistical analyses were performed in GraphPad Prism v.10.2, except for the moving-average and regression analyses, which were performed in R. Specific tests used for analysis can be found in the figure legends.

## Acknowledgements

We thank members of the Stowers lab for critical feedback on this manuscript. We would like to acknowledge Dr. Juan Manuel Urueña at the NSF BioPACIFIC MIP for providing training and support on the Optics11 Pavone microindenter. We also want to acknowledge Iosla Saoirse Finn at the University of California, Santa Barbara for assistance with image analysis. This work was supported by an American Cancer Society Catalyst award to R.S.S. (CAT-24-1379187-01-CAT). A.S. was supported by the California Institute of Regenerative Medicine (CIRM-12821) EDUC4 training grant. M.A.G-G. was supported by an American Heart Association Postdoctoral Fellowship (24POST1195931). M.P. and J.A.B. were supported by a grant from the National Institutes of Health, USA (NIH grant 1T32GM141846). The content is solely the responsibility of the authors and does not necessarily represent the official views of the National Institutes of Health. M.P. was also supported by the UC Santa Barbara Regents in Bioengineering fellowship. This work was supported by the BioPACIFIC Materials Innovation Platform of the National Science Foundation under Award No. DMR-1933487. The heparan sulfate monoclonal antibody developed by D.M. Fambrough at Johns Hopkins University and the chondroitin sulfate monoclonal antibody developed by W.M. Halfter at University of Pittsburgh were obtained from the Developmental Studies Hybridoma Bank, created by the NICHD of the NIH and maintained at The University of Iowa, Department of Biology, Iowa City, IA 52242. We acknowledge the use of the NRI-MCDB Microscopy Facility at the University of California, Santa Barbara.

## Author Contributions

Conceptualization, A.S., J.A.B., R.S.S.; methodology, A.S., J.A.B., M.P., R.S.S.; software, A.S.; validation, A.S., J.A.B., M.P., M.A.G-G.; formal analysis, A.S., J.A.B., M.P., M.A.G-G.; investigation, A.S., J.A.B., M.P.; resources, R.S.S.; writing - original draft, A.S., J.A.B., R.S.S.; writing - review and editing, A.S., J.A.B., M.P., M.A.G-G., R.S.S.; visualization, A.S., J.A.B., M.P.; supervision, R.S.S.; funding acquisition, R.S.S.

## Data Availability

Data are available from the corresponding author upon reasonable request.

## Supplementary information

**Supp. Fig. 1:**
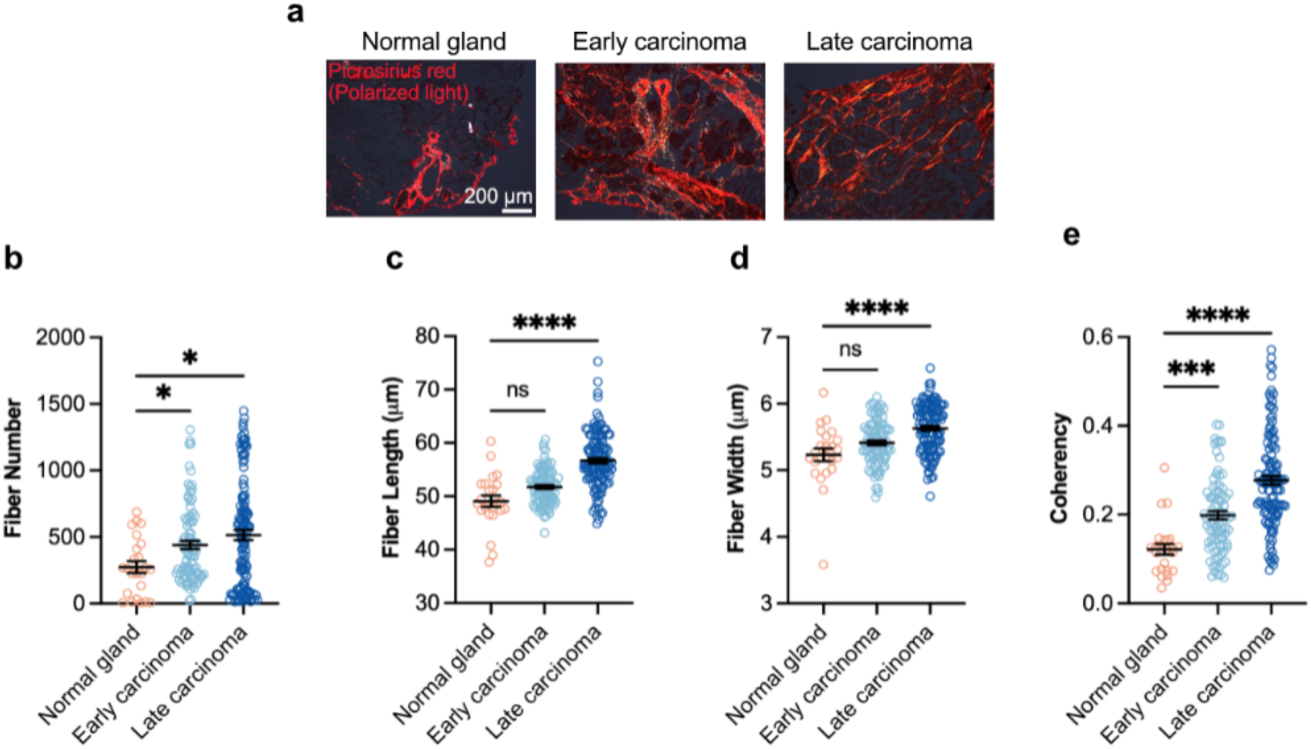
Tumor progression is associated with fibrillar collagen remodeling. **a**, Picrosirius red stained tissues visualized under polarized light showed thick, aligned fibrillar collagen bundles (in red) in early and late carcinoma tissues compared to the normal gland. **b,** Quantification of collagen fiber parameters showed a significant increase in the number of collagen fibers in late carcinoma samples compared to normal glands. **c,** Fiber length and **d,** fiber width were significantly higher in late carcinoma samples than in normal glands. **e,** Local fiber orientation in regions of interest (ROIs) was quantified using the coherency parameter, ranging from 0 (isotropic) to 1 (highly aligned), in the OrientationJ macro in ImageJ. Both early and late carcinoma samples had a significantly higher fiber coherency compared to normal glands. For **b-d,** n ≥ 4 images were acquired per tumor, with at least 3 independent tumors used per group, and each tumor derived from a different mouse. Each image represents one data point. For **e,** at least 3 ROIs were recorded per image, and averaged per image, with at least 3 independent tumors used per disease group, and each tumor derived from a different mouse. For **b-e,** mean ± s.e.m.; Kruskal-Wallis test followed by post-hoc Dunn’s multiple comparison test was performed. ∗p < 0.05; ∗∗p < 0.01; ∗∗∗p < 0.001; ∗∗∗∗p < 0.0001, not significant = ns.

**Supp. Fig. 2:**
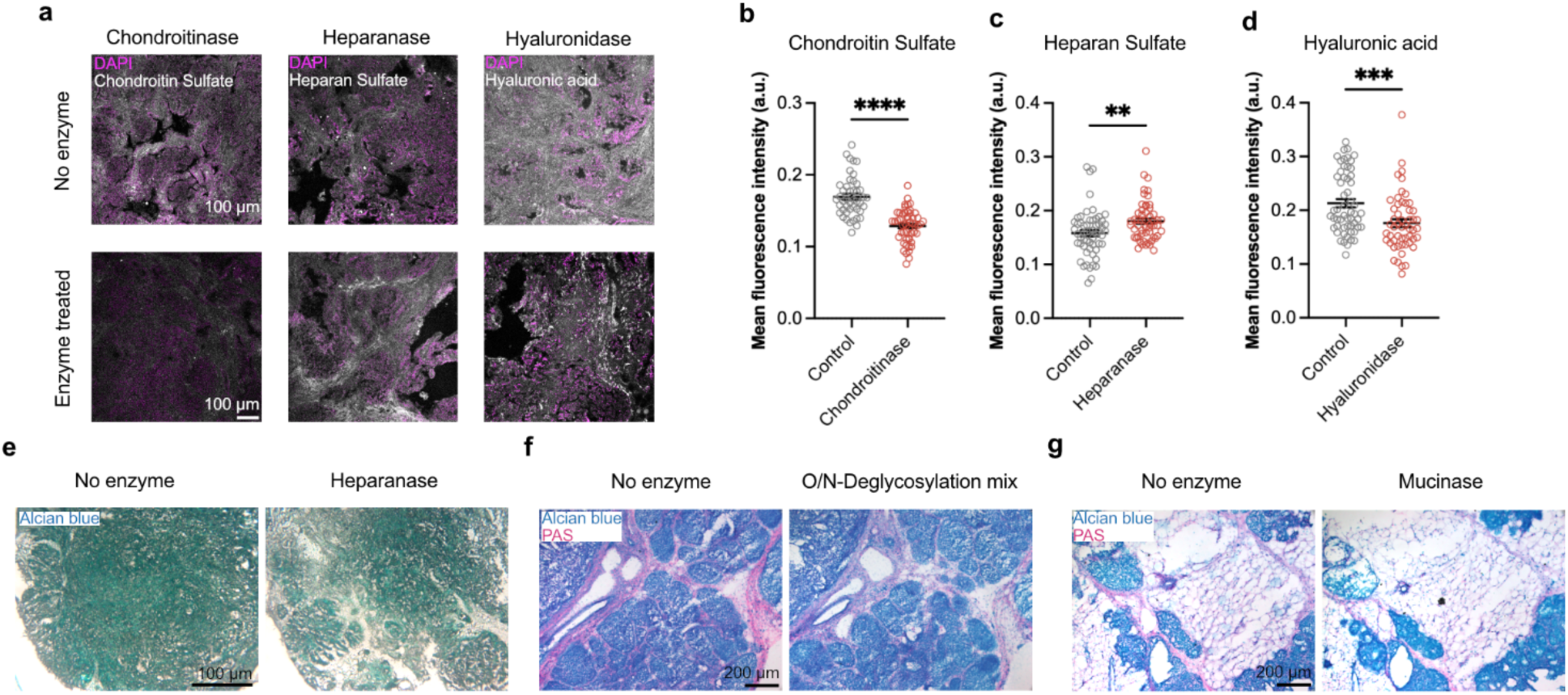
Enzymatic degradation of GAGs and glycoproteins results in reduced staining intensity. **a**, Late carcinoma tissue sections were stained for chondroitin sulfate, heparan sulfate, and hyaluronic acid after treatment with chondroitinase, heparinase, and hyaluronidase respectively. Buffer without enzyme was used as the control. Mean intensity of **b,** chondroitin sulfate and **d,** hyaluronic acid was significantly reduced upon treatment. **c,** Heparan sulfate instead showed a significantly increased intensity upon treatment with heparinase, possibly due to increased access to the epitope located on the protein backbone of the proteoglycan after treatment. **e,** Alcian blue staining was used to verify reduced GAG abundance after treatment of tissue section with heparinase. **f,** PAS stain (pink), used to visualize glycoproteins, showed reduced staining upon treatment of tissue section with the O/N-deglycosylation mix. Another section from the same tumor treated with a buffer without enzyme was used as a negative control. **g,** PAS stain used to visualize degradation of mucins after treatment of a late carcinoma tissue section with StcE mucinase, also showed reduced staining compared to the no enzyme control. For **b-d,** n ≥ 15 images per tissue, with at least 3 independent tissue samples per group, and each tissue from a different mouse (mean ± s.e.m.; Mann-Whitney test was performed). ∗p < 0.05; ∗∗p < 0.01; ∗∗∗p < 0.001; ∗∗∗∗p < 0.0001, not significant = ns.

**Supp. Fig. 3:**
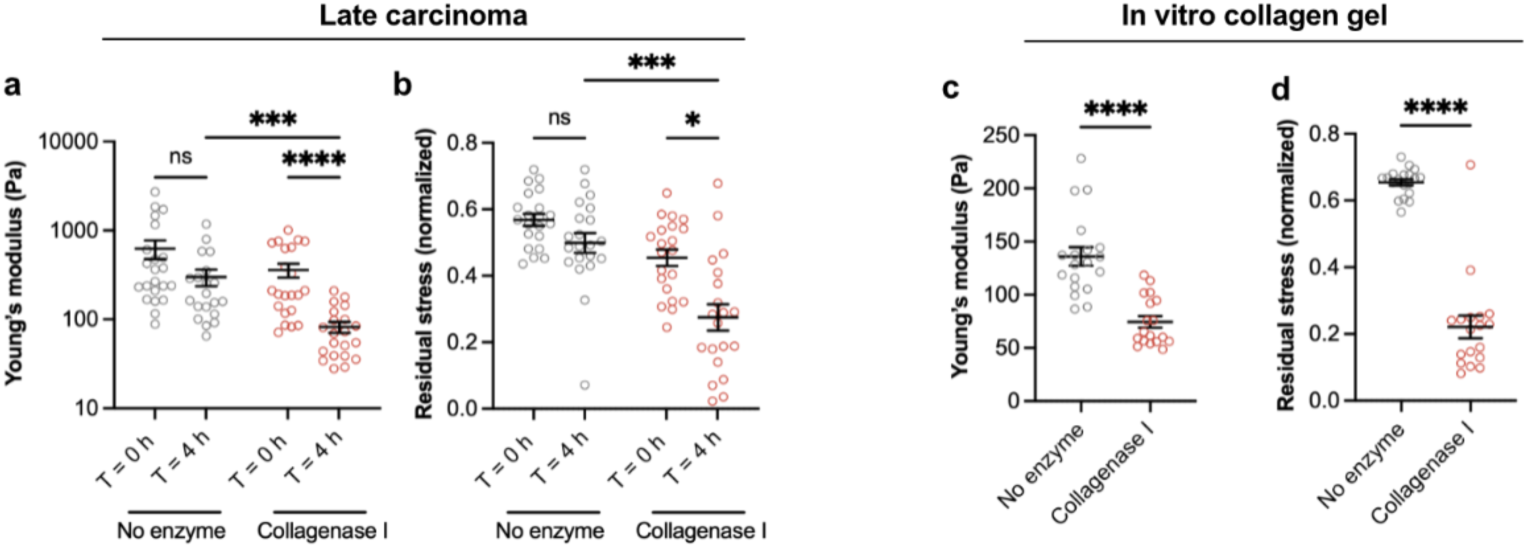
Late carcinoma tissue treated with collagenase showed reduced Young’s modulus and increased viscoelasticity. **a**, Late carcinoma samples were incubated with 0.5 mg/ml collagenase for 4 h and indented before and after treatment. Samples showed a significantly reduced Young’s modulus and also **b,** significantly reduced residual stress (200 s), indicating increased stress relaxation, compared to the no enzyme controls. n = 5 indentations per tissue, with 3 independent tissue samples per treatment group, and each tissue derived from a different mouse (mean ± s.e.m.; Kruskal-Wallis test followed by post-hoc Dunn’s multiple comparisons test). 2 mg/ml collagen I gels treated with collagenase for 1 h also showed a significant reduction in **c,** Young’s modulus and **d,** residual stress. n = 6 indentations per gel, with 3 independent gels per treatment group (mean ± s.e.m.; Mann-Whitney test). ∗p < 0.05; ∗∗p < 0.01; ∗∗∗p < 0.001; ∗∗∗∗p < 0.0001, not significant = ns.

**Supp. Fig. 4:**
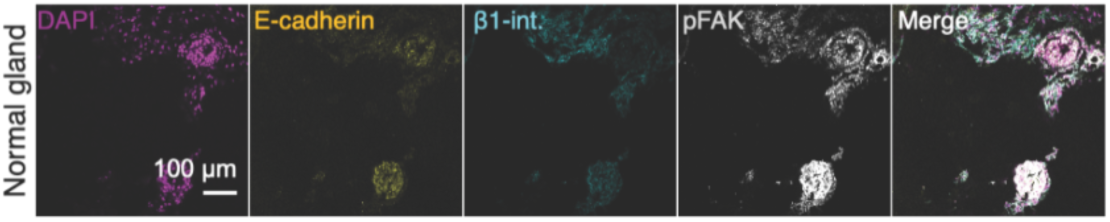
Normal mouse glands show high pFAK expression. Representative images of normal mouse gland tissue stained for DAPI, E-cadherin, β1-integrin, and FAK phosphorylated at Tyr397 (pFAK). Scale bars, 100 μm.

**Supp. Fig. 5:**
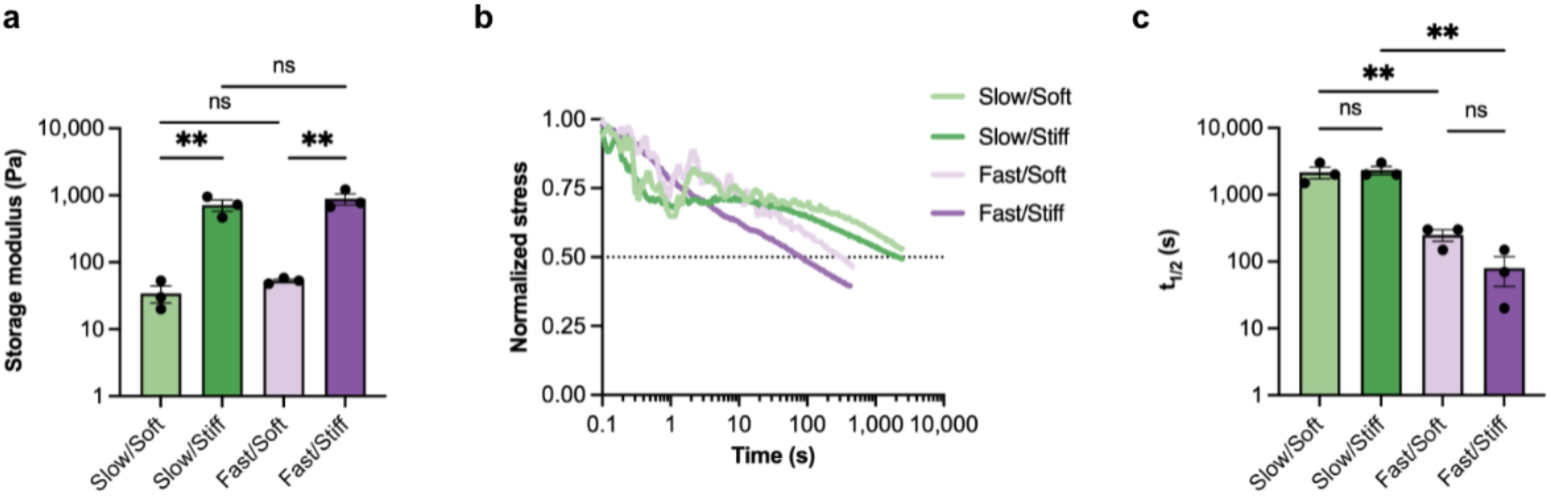
Alginate-rBM IPNs allow for independent tuning of stiffness and viscoelasticity. **a**, Storage modulus of gels measured at 0.5% strain and 1 Hz was significantly higher in stiff/slow and stiff/fast groups, mimicking the stiffness of carcinoma samples, than in soft/slow and soft/fast groups, mimicking the normal gland stiffness. **b,** Stress relaxation profiles of IPNs measured at a 10% constant strain. **c,** Soft/slow and stiff/slow groups had a significantly greater stress relaxation time, t1/2, to reach half of the maximum stress, than the soft/fast and stiff/fast groups. For **a, c,** n = 3 gels per group (mean ± s.e.m.; one-way ANOVA test followed by Holm-Sidak’s multiple comparison test was performed). ∗p < 0.05; ∗∗p < 0.01; ∗∗∗p < 0.001; ∗∗∗∗p < 0.0001, not significant = ns.

